# Population receptive field and connectivity properties of the early visual cortex in human albinism

**DOI:** 10.1101/627265

**Authors:** Khazar Ahmadi, Anne Herbik, Markus Wagner, Martin Kanowski, Hagen Thieme, Michael B. Hoffmann

## Abstract

In albinism, the pathological decussation of the temporal retinal afferents at the optic chiasm leads to superimposed representations of opposing hemifields in the visual cortex. Here, we assessed the equivalence of the two representations and the cortico-cortical connectivity of the early visual areas. Applying fMRI-based population receptive field (pRF)-mapping (both hemifield and bilateral mapping) and connective field (CF)-modeling, we investigated the early visual cortex in 6 albinotic participants and 4 controls. In albinism, superimposed retinotopic representations of the contra- and ipsilateral visual hemifield were observed on the hemisphere contralateral to the stimulated eye. This was confirmed by the observation of bilateral pRFs during bilateral mapping. Hemifield mapping revealed similar pRF-sizes for both hemifield representations throughout V1 to V3. The typical increase of V1-sampling extent for V3 compared to V2 was not found for the albinotic participants. The similarity of the pRF-sizes for opposing visual hemifield representations highlights the equivalence of the two maps in the early visual cortex. The altered V1-sampling extent in V3 indicates the adaptation of cortico-cortical connections to the abnormal input of the visual cortex. These findings thus suggest that conservative developmental mechanisms are complemented by alterations of the extrastriate cortico-cortical connectivity.

**Highlights:**

- pRF mapping confirms cortical overlay of opposing visual hemifields in albinism.
- Equivalent information processing of both hemifields is indicated by similar pRF sizes.
- CF modeling indicates changes to the cortico-cortical connections at the level of V3.

## 1. Introduction

Albinism is associated with misrouted optic nerves, which leads to sizable abnormal retinotopic organization in the visual cortex (Guillery, 1986; Hoffmann and Dumoulin, 2015). Typically, nasal retinal fibers cross the midline at the optic chiasm and terminate in the contralateral hemisphere, while fibers originating in temporal retina stay uncrossed and project to the ipsilateral hemisphere. The line of decussation thus coincides with the vertical meridian through the fovea. As a result of this projection scheme, each hemisphere receives binocular input from the contralateral visual field. This input is initially segregated into interdigitated ocular dominance domains in V1, but converges in extrastriate areas to yield binocular visual function and stereopsis (Hubel and Wiesel, 1968; Parker et al., 2016). The normal projection of the retinal fibers is substantially altered in albinism i. e. the line of decussation is shifted toward the temporal retina such that a greater extent of temporal retinal axons project contralaterally (Apkarian et al., 1983; Creel, 1971; Guillery et al., 1975). As a consequence, each hemisphere receives predominantly monocular input from the ipsilateral visual field in addition to the normal input from the contralateral visual field (Schmitz et al., 2004; von dem Hagen et al., 2008). This results in superimposed monocular retinotopic maps of opposing hemifields (Hoffmann et al., 2003; Kaule et al., 2014), which disrupts the integration of input from both eyes and subsequently binocular and stereo-vision (Hoffmann and Dumoulin, 2015). When inspected at higher spatial resolution, as demonstrated electrophysiologically in an albino green monkey (Guillery et al., 1984), these superimposed maps form hemifield dominance domains that are reminiscent of ocular dominance domains in a normal visual system. Despite the substantial aberrant input to the visual cortex, major aspects of visual function are preserved (Eick et al., 2019; Hoffmann et al., 2017; Hoffmann and Dumoulin, 2015; Klemen et al., 2012; Wolynski et al., 2010). This is taken as evidence for adaptive mechanisms that make the erroneous visual input available for perception and highlights the importance of albinism as a powerful model to study the foundation of visual pathway formation and the scope of plasticity in humans.

The aim of the present study was to determine the consequences of atypical visual projections on population receptive field (pRF; Dumoulin & Wandell, 2008) and cortical connective field (CF; Haak et al. 2012) properties in albinism. We confirm superimposed retinotopic representations of opposing visual hemifields in albinism and report similar pRF sizes and hence equivalent processing for both hemifield representations. Furthermore, we observe changes to the extrastriate cortico-cortical connections in albinism at the level of V3. Our results thus provide independent evidence for a lack of large-scale reorganization and suggest that alterations of the intra-cortical and cortico-cortical connectivity of the visual system compensate for the substantial projection abnormality of the optic nerves in albinism.

## 2. Methods

### 2.1 Participants

Six albinotic participants (mean age = 35, range = 18 – 60 years; 3 females) were recruited for this study. In an ophthalmological examination, the typical symptoms of albinism were identified (iris transillumination, foveal hypoplasia, hypopigmentation fundus) and the decussation abnormality was confirmed with misrouting-VEPs according to the procedure described previously (Hoffmann et al., 2015). Absence of stereo-vision was verified using Lang I, Titmus, and TNO-tests. Monocular best-corrected decimal visual acuities were assessed and horizontal fixation stability was determined with a fundus-controlled measurement (MP-1 microperimeter, Nidek, Padova, Italy). Detailed characteristics of the albinotic participants are reported in Table 1. In addition, four controls (mean age = 31, range = 25 – 49 years; 2 females) with normal visual acuity, normal stereo-vision, and no history of ophthalmological or neurological disorders participated in this study. All participants gave their informed written consent. The study was approved by the ethics committee of the University of Magdeburg and the procedure adhered to the tenets of the declaration of Helsinki.

**Table 1.**
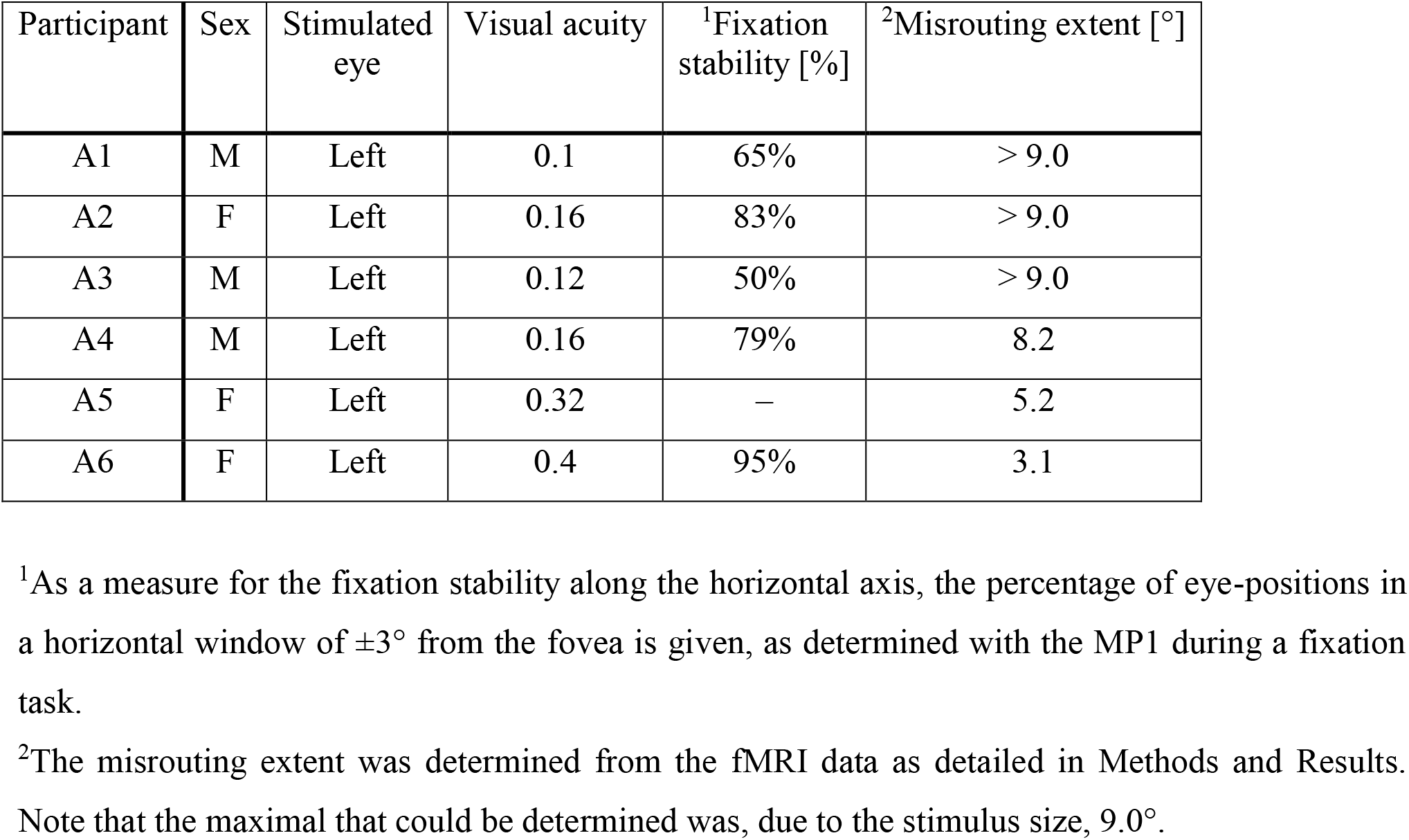
Characteristics of the albinotic participants

| Participant | Sex | Stimulated eye | Visual acuity | <sup>1</sup>Fixation stability [%] | <sup>2</sup>Misrouting extent [°] |
| --- | --- | --- | --- | --- | --- |
| A1 | M | Left | 0.1 | 65% | > 9.0 |
| A2 | F | Left | 0.16 | 83% | > 9.0 |
| A3 | M | Left | 0.12 | 50% | > 9.0 |
| A4 | M | Left | 0.16 | 79% | 8.2 |
| A5 | F | Left | 0.32 | – | 5.2 |
| A6 | F | Left | 0.4 | 95% | 3.1 |

<sup>1</sup>As a measure for the fixation stability along the horizontal axis, the percentage of eye-positions in a horizontal window of $\pm 3^\circ$ from the fovea is given, as determined with the MP1 during a fixation task.
<sup>2</sup>The misrouting extent was determined from the fMRI data as detailed in Methods and Results. Note that the maximal that could be determined was, due to the stimulus size, $9.0^\circ$ .

### 2.2 MRI acquisition

Functional T2*-weighted echo-planar volumes were acquired using a 3T Magnetom Prisma scanner with the 64 channel head coil (Siemens Healthineers, Erlangen, Germany). An Original Pillow Junior (Tempur-Pedic) was placed on the base of the coil surrounding the sides and the back of the head for a good balance between comfort and reduction of head motion. The data were obtained at an isotropic resolution of 2.5 x 2.5 x 2.5 mm^3^ with 54 axial slices covering the whole brain (TR | TE = 1500 ms | 30 ms, flip angle: 70°, FOV = 210 mm, multi-band and in-plane acceleration factors = 2). Each functional scan was 168 time frames (252 s) in duration. A total of 9 functional scans were acquired in a single session [three repetitions per experimental configuration (see below, Visual stimulation)]. Additionally, a T1-weighted anatomical volume was collected at the beginning of each session (MPRAGE; voxel size = 0.9 x 0.9 x 0.9 mm^3^, TR | TI | TE = 2600 ms | 1100 ms | 4.46 ms, and flip angle = 7°).

### 2.3 Visual stimulation

Drifting bar apertures (9.5° in radius), exposing a moving high-contrast checkerboard pattern (Dumoulin and Wandell, 2008) were displayed at four directions i.e. upward, downward, left and right. The bar moved across the stimulus window in 20 evenly spaced steps and its width subtended 1/4th of the stimulus radius. Each pass of the bar lasted for 30 s, followed by a mean luminance block (zero contrast) of 30 s. The stimuli were generated in Matlab (Mathworks, Natick, MA, USA) using the Psychtoolbox (Brainard, 1997; Pelli, 1997) and projected onto a screen with a resolution of 1140 x 780 pixels at the magnet bore. Participants viewed the screen monocularly at the distance of 35 cm via an angled mirror and their dominant eye was stimulated under three experimental configurations: (i) bilateral, (ii) left, and (iii) right hemifield stimulation (Ahmadi et al., 2018). They were required to fixate a centered dot and to report color changes between red and green via button press.

### 2.4 Data preprocessing and analysis

The T1-weighted anatomical volume was automatically segmented using FreeSurfer (https://surfer.nmr.mgh.harvard.edu). The cortical surface was reconstructed at the white/gray matter boundary and rendered as a smoothed 3D mesh (Wandell et al., 2000). FSL (https://www.fmrib.ox.ac.uk/fsl) was used for the correction of motion artefacts in the functional data. Motion-corrected data for each experimental configuration were then averaged together for every participant to increase the signal-to-noise ratio (SNR). Subsequently, the functional data were aligned to the anatomical volume using a combination of Vistasoft tools (https://github.com/vistalab/vistasoft) and Kendrick Kay’s alignment toolbox (https://github.com/kendrickkay/alignvolumedata). All further analyses, including the estimation of pRF and CF properties, the delineation of the visual areas and the visualization on the smoothed mesh surface were performed in Vistasoft. The pRF sizes and positions were estimated from the fMRI data and visual stimulus position time course. The BOLD (blood oxygen level dependent) response of each voxel was predicted using a circular 2D-Gaussian model of the neuronal populations defined by three stimulus-referred parameters i.e. x_0_, y_0_, σ where x_0_ and y_0_ are the coordinates of the receptive field center and σ is it’s spread (Dumoulin and Wandell, 2008; Fracasso et al., 2016; Harvey and Dumoulin, 2011). The predicted BOLD signal was calculated by convolution of the stimulus sequence for the respective pRF-model and its three parameters with the canonical hemodynamic response function (Friston et al., 1998). The optimal pRF parameters were found by minimizing the residual sum of squared errors (RSS) between the predicted and observed BOLD time-course. Only voxels were retained whose explained variance exceeded a threshold of 15%. To assess the presence of bilateral pRFs in V1 to V3, we extended the conventional pRF model in analogy to previous studies (Ahmadi et al., 2018; Fracasso et al., 2016; Hoffmann et al., 2012). As such, we compared the conventional pRF model with mirror-pRF models across the (i) vertical meridian and (ii) horizontal meridian, here termed as mirror-pRF models across VM and HM, respectively. While the conventional pRF model consists of a single circularly symmetric 2D Gaussian, the mirror-pRF models comprise two 2D Gaussians that are mirrored across the vertical or horizontal meridians. Because all parameters of the two Gaussians are linked to each other, mirror-pRF models have the same degrees of freedom as the conventional pRF model. Consequently, the model performance can be compared directly. Unlike the conventional pRF model, mirror-pRF models predict that each cortical location responds to two distinct regions in the visual field.

The pRF model is prone to biased estimates of the receptive fields for visual stimuli that comprise masks, as opposed to full-field stimulation within a circular aperture. This particularly affects pRFs that are located at the edge of the stimulus space (Binda et al., 2013; Lee et al., 2013; Papanikolaou et al., 2015). To avoid this problem, we excluded the representations of the vertical meridian (coinciding with the edge of the hemifield stimuli) from each region of interest (ROI) for every participant. In addition, the ROIs were restricted to the regions with the overlap of both hemifields for the albinotic participants.

The CF parameters were estimated from the fMRI time-series, using CF modeling that predicts the neuronal activity in one brain area with reference to aggregate activity in another area (Haak et al., 2013). Briefly, the BOLD response in each voxel of a target ROI i.e. V2 or V3 was predicted with a symmetrical, circular 2D Gaussian CF model folded to follow the cortical surface of the source ROI i.e. V1. The CF model was defined by two parameters, namely, Gaussian position and spread across the cortical surface. The optimal CF parameters were determined by minimizing the RSS between the predicted, and the observed time-series. For this purpose, many fMRI time-series predictions were generated by changing the CF positions across all voxel positions and Gaussian spread values on the surface of the source ROI. As for the pRF-modelling, only model fits were selected whose explained variance exceeded a threshold of 15%. We obtained V1 sampling extent in V2 and V3 by adjusting the V1-referred CF size in those areas for pRF laterality i.e. the extent to which a pRF overlaps with the ipsilateral visual field (Haak et al., 2013).

### 2.5 Statistical analysis

One-sample t-tests were performed to compare (i) the difference of the pRF sizes of V1 to V3 between the two stimulated hemifields and (ii) the difference of the V1-sampling extent between V3 and V2 in each of the albinotic and control groups. When applicable, multiple comparisons were corrected using the Bonferroni-Holm procedure (Holm, 1979) and the adjusted alpha level for each comparison was denoted as (p_α_). Additionally, a linear regression model was used to assess the dependence of the V1-sampling extent on eccentricity.

## 3. Results

We investigated the functional properties of the early visual areas (V1, V2, and V3) of the albinotic participants in two steps. Firstly, based on bilateral and hemifield pRF-mapping, we detailed the pRF properties of the visual field maps. Secondly, we applied CF-modeling to determine V1-sampling extent in V2 and V3.

### 3.1 Visual field maps and pRF properties in albinism

The visual field map properties obtained for pRF mapping are given in Figure 1. Here the eccentricity and polar angle maps obtained for hemifield pRF-mapping are juxtaposed for a control and two individuals with albinism (all stimulated via the left eye). The left hemifield was represented as an orderly eccentricity and polar angle map on the contralateral, i.e. right hemisphere, in both control and albinism, confirming the normal projection of the nasal retina for all conditions. In contrast, misrouting of the temporal retinal fibers (Hoffmann and Dumoulin, 2015) was evident for the representation of the right hemifield in both albinotic participants. Here orderly eccentricity and polar angle maps were found on the right hemisphere, i.e., ipsilateral to the stimulated hemifield. In one of the depicted individuals with albinism (Figure 1 B) this abnormality was extensive, indicating a larger shift of the line of decussation into the temporal retina than for the other individual (Figure 1 C). This is in accordance with the well-known variability of misrouting in albinism (Hoffmann et al., 2005, 2003; von dem Hagen et al., 2007). In Table 1 the extent of misrouting is provided, as determined from the mean eccentricity value of an ROI covering the anterior activated margin of the abnormal representation of the horizontal meridian in the right V1.

**Figure 1:**
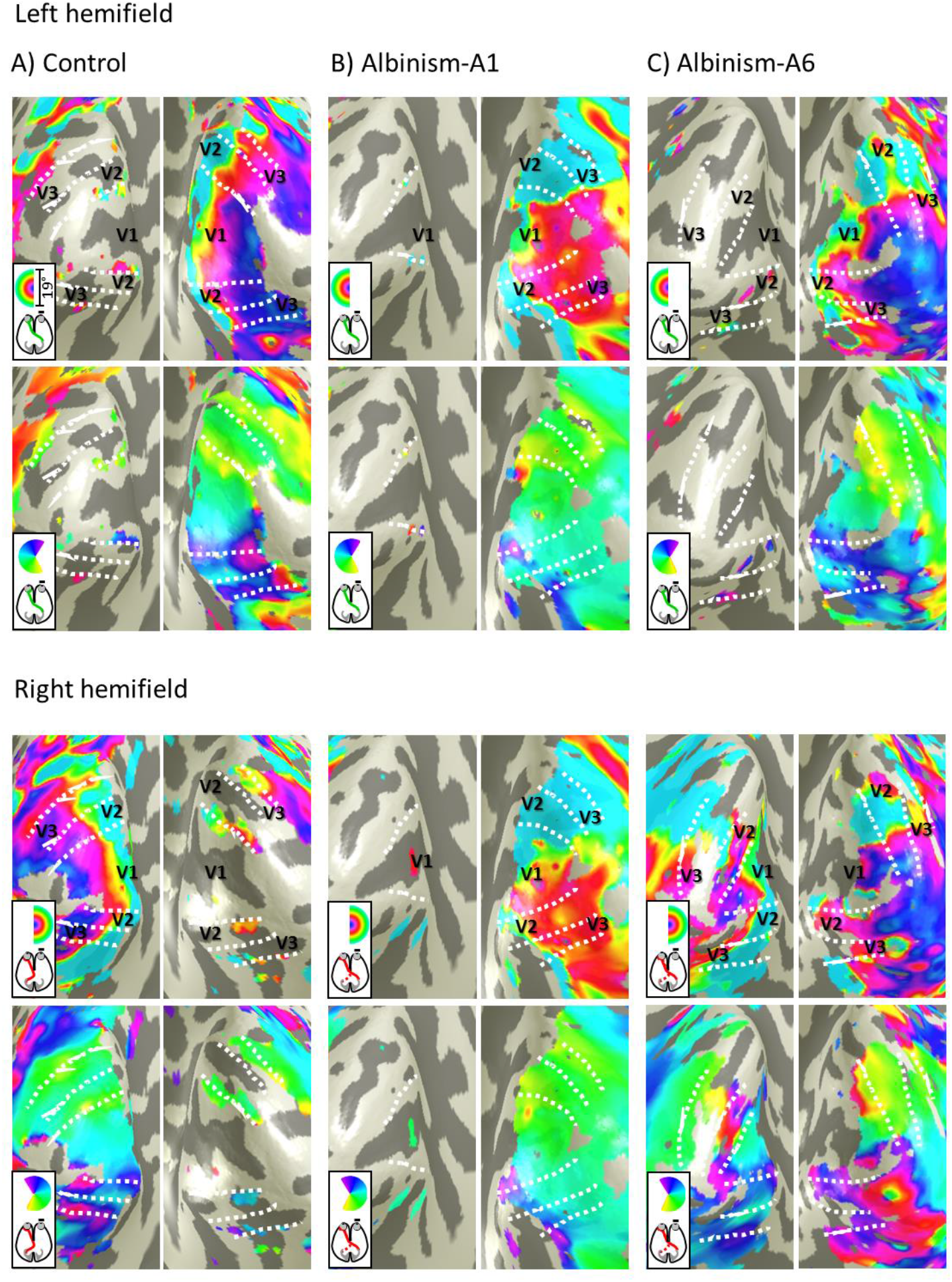
Hemifield pRF-mapping. Eccentricity and polar angle maps (top and bottom rows in the two panels, respectively) are shown on the inflated occipital cortex under the left (nasal retina; top panel) and right (temporal retina; bottom panel) hemifield stimulation conditions. **(A)** In the control, stimulation of each visual hemifield elicits orderly eccentricity and polar angle maps predominantly on the hemisphere contralateral to the stimulated hemifield. Only, residual representations of the vertical meridian and fovea are observed on the ipsilateral hemisphere, as reported previously (Hoffmann et al., 2003; Tootell et al., 1998). **(B & C)** In contrast, in albinism there is, in addition, to the representation of the contralateral (left) visual hemifield, a representation of the ipsilateral (right) hemifield on the right hemisphere, i.e. contralateral to the stimulated eye. While this is extensive for A1, it is smaller for A6 (see Table 1), where, as a consequence, a residual normal right hemifield representation is evident on the left hemisphere. Note that the slight deviation of the eccentricity color key for A1 is likely due to severe foveal hypoplasia.

All in all, the above pRF-hemifield mapping findings demonstrate the mirror-symmetrical retinotopic cortical overlay of normal and abnormal representations of the contralateral and ipsilateral visual hemifield respectively in albinism (Hoffmann et al., 2003; Kaule et al., 2014). This was independently confirmed by the bilateral pRF-mapping data: In analogy to previous studies on FHONDA syndrome (<u>f</u>oveal <u>h</u>ypoplasia, <u>o</u>ptic <u>n</u>erve <u>d</u>ecussation defects and <u>a</u>nterior segment dysgenesis), hemihydranencephaly, and achiasma (Ahmadi et al., 2018; Fracasso et al., 2016; Hoffmann et al., 2012), the goodness of fit, i.e. variance explained (VE), was compared between (i) the conventional single-pRF model and the mirror-pRF models, (ii) across VM, expected to reflect the mirror-symmetrical overlay of opposing hemifields in albinism, and (iii) across HM, as a reference model. In fact, the mirror-pRF model across VM performed worst in the controls, while it performed better in the two albinotic participants with below average misrouting (8° as determined in Hoffmann et al., 2005), i.e., misrouting (MR) < 8°, and best in those with above average misrouting, i.e., MR > 8° (see Figure 2). Importantly, this does not only apply to V1, but also to V2 and V3, and thus confirms overlaid representations in these three early visual areas in albinism.

**Figure 2:**
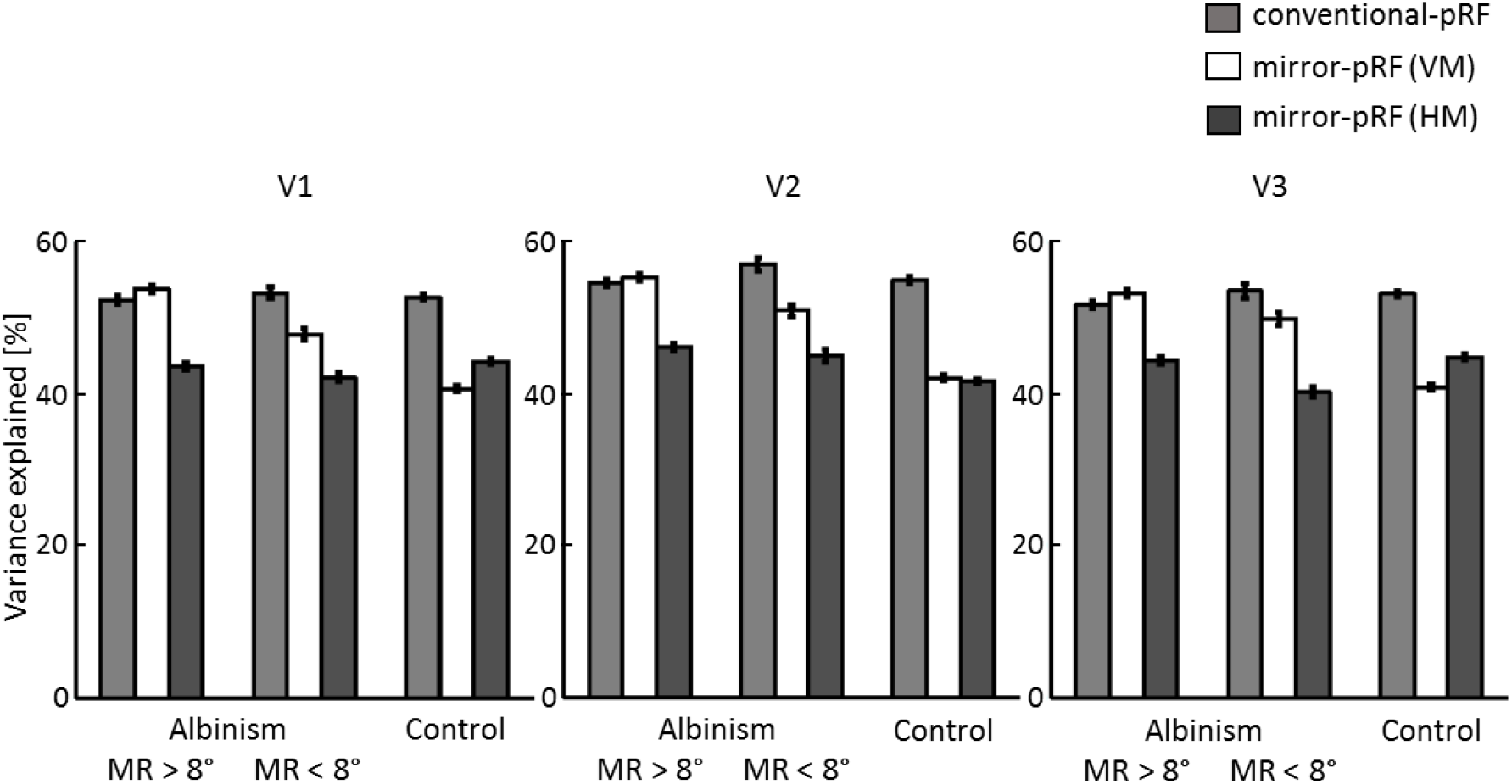
Comparison of the explanatory power of different pRF-models across participant groups and visual areas. The mean VE ± SEM is depicted for the conventional pRF model (light gray bars), mirror-pRF models across VM (white bars) and HM (dark gray bars) in V1 to V3. The mirror-pRF model across VM performs worst in the controls, better in albinism with misrouting extent (MR) < 8° and best in albinism with MR > 8° in all three visual areas.

We examined the results for the hemifield stimulation conditions further, to test the functional equivalence of the two superimposed hemifield representations in albinism. Specifically, we compared the pRF sizes for both hemifield representations. The average pRF sizes of the albinotic participants increased as a function of eccentricity and through visual hierarchy for both hemifield representations, albeit more scattered for the right hemifield representation (Figure 3 A). Note that despite the overall similarity of the pRF size vs eccentricity relationship between the albinotic and control participants, the average pRF sizes in albinotic individuals exceeded those of controls, likely due to fixation instabilities (Table 1). Subsequently, to assess the equivalence of the pRF sizes for the two hemifield representations, we subtracted the average pRF sizes of each area for the left hemifield from the corresponding average pRF sizes of the right hemifield representation (Figure 3 B). There was no significant difference of the pRF sizes of V1, V2, and V3 between the two hemifield representations on the same hemisphere in albinism (t (5) = −1.46, p_0.025_ = 0.20; t (5) = – 2.84, p_0.017_ = 0.04; t (5) = 0.17, p_0.05_ = 0.87, respectively), only a small non-significant trend (< 0.5°) was observed for larger pRF sizes in V1 and V2 in albinism for the abnormal, i.e. right, hemifield. Similarly, in controls no significant pRF-size differences were evident in V1 to V3 for the two hemifields represented on separate hemispheres (t (3) = 1.77, p_0.025_ = 0.17; t (3) = −2.02, p_0.017_ = 0.14; t (3) = 1.66, p_0.05_ = 0.20, respectively). Taken together, in albinism, V1, V2, and V3 comprise functionally equivalent superimposed maps of both the contra- and ipsilateral visual hemifield.

**Figure 3:**
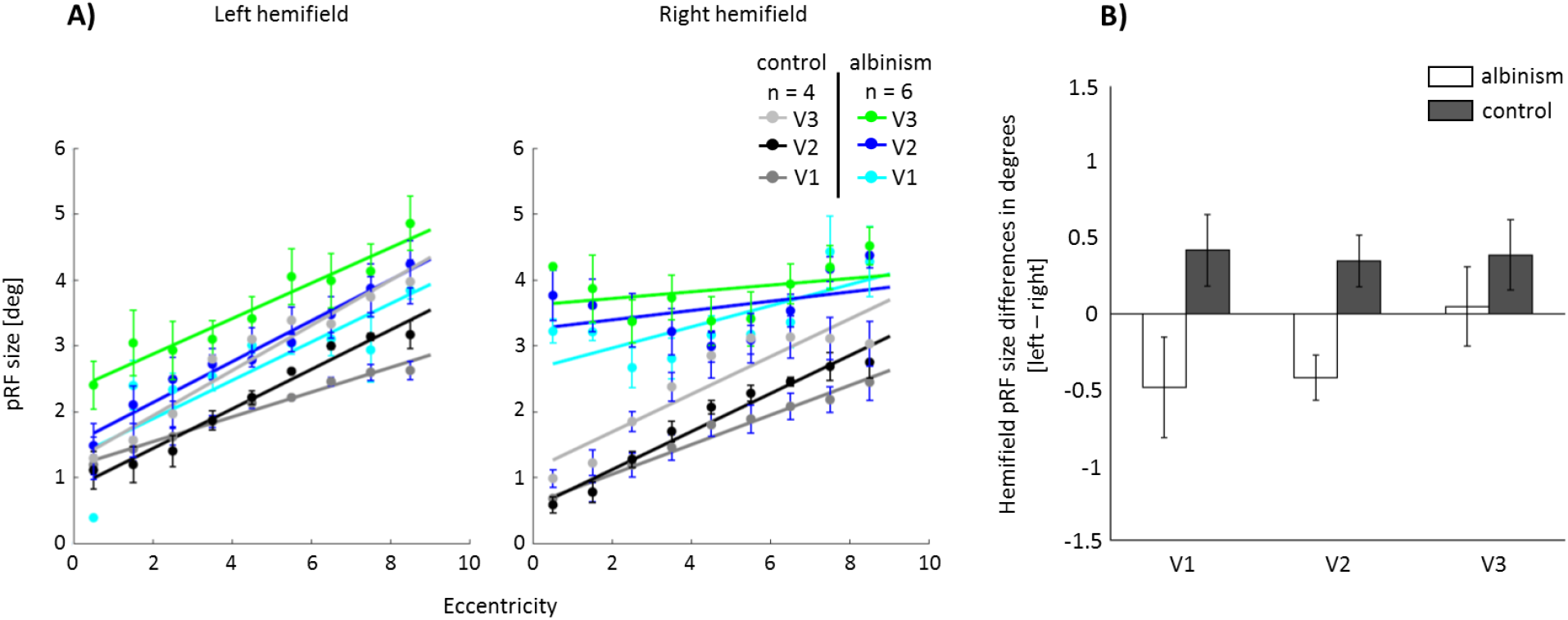
Eccentricity dependence and differences of pRF size between the left and right hemifield representations across participant groups and visual areas. **(A)** Similar to controls, the average pRF sizes of all albinotic participants increase as a function of eccentricity and through visual hierarchy for both hemifield stimulations. **(B)** No significant pRF-size differences were evident neither in the albinotic group nor in the controls. The difference of the pRF sizes between the two hemifields was first calculated in every eccentricity for each participant and subsequently grouped across participants. The bars represent the mean pRF size difference in each group and error bars indicate SEM.

### 3.2 CF-properties in albinism

Generally, the superimposed maps from opposing hemifields in V1 in albinism are taken as evidence for largely conservative, i.e., stable, geniculo-striate connections and a reassignment of ocular dominance domains to hemifield domains. Likewise, the propagation of this mapping-scheme to V2 and V3 suggests largely conservative cortico-cortical connections (Hoffmann and Dumoulin, 2015). However, due to the binocular nature of cells in extrastriate areas of the neuro-typical visual system, beyond V1 no simple reassignment of ocular dominance domains is available as a mechanism to accommodate the extra-map of the ipsilateral visual field in albinism. This could reflect on the V1-sampling extent in these areas. We applied CF-modeling to investigate the cortico-cortical functional connectivity profiles in albinism for the same ROIs also used for the pRF assessments and compared the V1-sampling extent of V2 and V3 averaged across the participant groups and the two hemifield representations. For the controls, the average V1-sampling extent in V2 and V3 was, in accordance with previous reports (Haak et al., 2013), roughly constant across eccentricity (R^2^ = 0.10, p = 0.43, and R^2^ = 0.28, p = 0.17, respectively), and increasing through the visual hierarchy. A similar independence from eccentricity was evident for albinism V2 (R^2^ = 0.13, p = 0.37), while a potential dependence on eccentricity was observed for V3 (R^2^ = 0.93, p < 0.00001; see Figure 4 A). This deviation of V3 from the neuro-typical condition was further supported by the comparison of the difference of V1-sampling-extent observed for V2 and V3 (Figure 4 B). While it increased from V2 to V3 in controls (t (3) = 3.24, p = 0.04) as reported previously (Gravel et al., 2014; Haak et al., 2013), no increase was evident for albinism (t (5) =1.1, p =0.32). Taken together, these findings indicate a largely unaltered V1-V2 connectivity in albinism, but suggest an alteration of the functional connectivity for V3.

**Figure 4:**
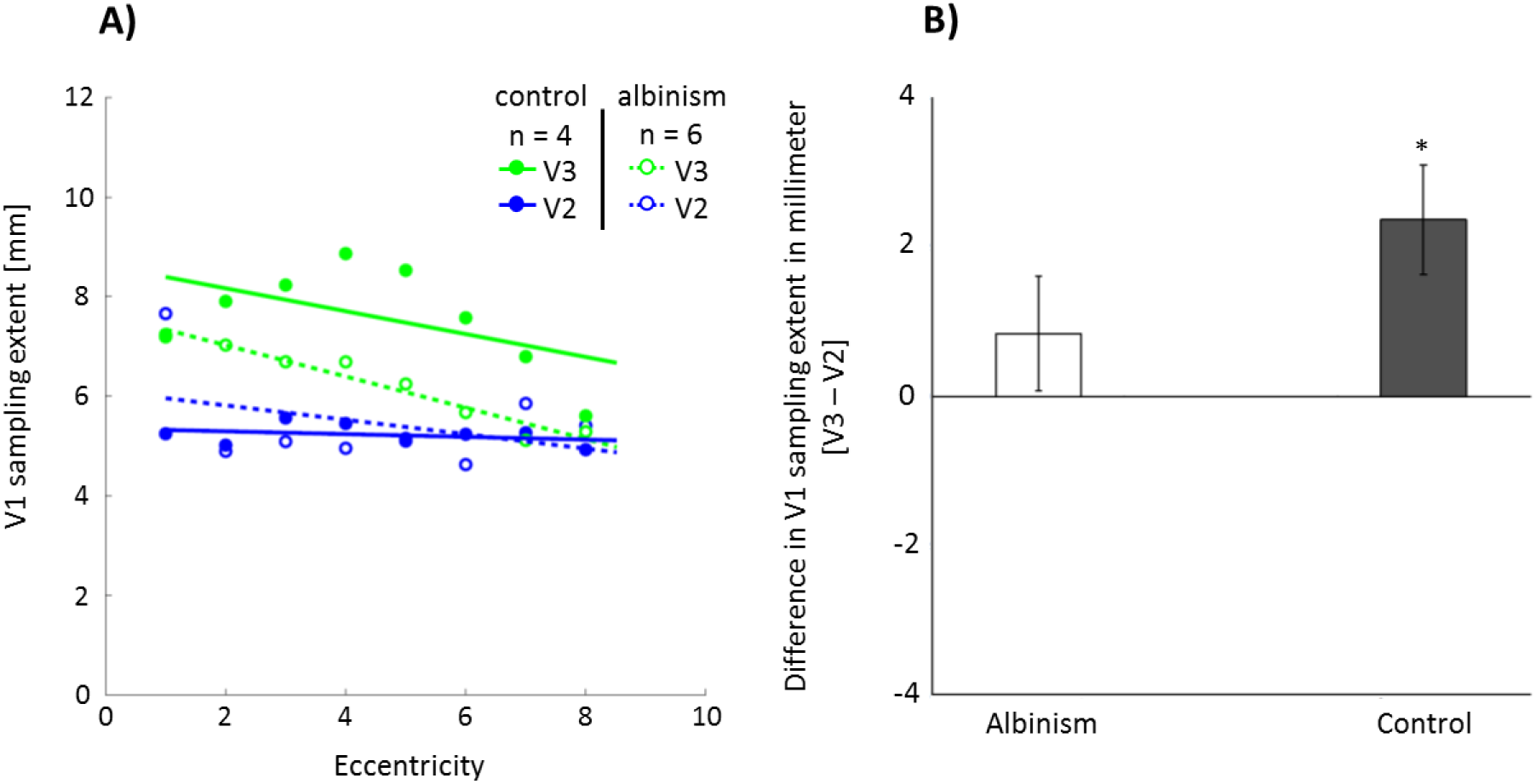
Comparison of V1-sampling extent in V2 and V3 across participant groups. **(A)** Eccentricity dependence of V1-sampling extent grouped across participants and stimulated hemifields. Akin to the controls, the average V1-sampling extent in V2 remains relatively constant across eccentricity in albinism. However, there is a trend for a decreasing eccentricity dependence of the average V1-sampling extent in V3. Eccentricity is binned in intervals of 1°. Each dot indicates the mean size of V1-sampling extent for every eccentricity bin, and solid lines demonstrate the linear fits for the dots. **(B)** The difference of V1-sampling extent between V3 and V2. The difference of V1-sampling extent in V3 and V2 was averaged across eccentricity and subsequently across hemifields in albinotic participants (white bar) and across hemispheres in controls (gray bar). The bars and error bars indicate the mean difference in V1 sampling extent ± SEM. While in controls the mean difference is significant and exceeds 2 mm, in albinotic participants this difference (0.8 mm) does not reach significance, indicating alterations in the functional connectivity between V1 and V3.

## 4. Discussion

We demonstrate that in albinism the superimposed maps of opposing visual hemifields in V1, V2, and V3 have similar pRF-sizes and that the cortico-cortical connectivity, as reflected by the V1-sampling extent, appears to be unaltered for V2, but altered for V3. This provides novel insights into the interplay of stability and plasticity supporting visual function in congenital visual pathway abnormalities.

### 4.1 pRF-mapping demonstrates equivalent superimposed retinotopic maps of opposing hemifields in albinism

We used pRF-mapping to detail the cortical organization in V1, V2 and V3. In accordance with previous findings, we demonstrated that the extent of the projection abnormality in albinism varies between individuals (Creel et al., 1981; Hoffmann et al., 2005; von dem Hagen et al., 2007) and that the abnormal input is mapped as a retinotopic overlay onto the normal input (Hoffmann et al., 2003; Kaule et al., 2014). As a consequence, mirror-symmetrical visual field positions are represented on similar cortical regions (Hoffmann and Dumoulin, 2015). In fact, voxels comprising these two hemifields can be modeled with bilateral receptive fields, as demonstrated in the present study for albinism and earlier for FHONDA, hemihydranencephaly and achiasma (Ahmadi et al., 2018; Fracasso et al., 2016; Hoffmann et al., 2012). This prompts the question of whether both representations are processed in the same manner. Unequal pRF-sizes for both representations would serve as an indicator of hemifield-specific processing differences. Remarkably, pRF sizes were equal for both hemifields, which provides physiological support for previous psychophysical reports on equivalent visual perception in both hemifields in albinism (Hoffmann et al., 2007; Klemen et al., 2012). These studies demonstrated similar sensitivities for visual perception mediated via the nasal or the, abnormally projecting, temporal retina, and a lack of cross-talk of information between the two hemifields. Thus, converging evidence is provided that both the contralateral and the additional ipsilateral hemifield representations in the early visual cortex are processed in a similar manner and independently of each other.

### 4.2 Conservative geniculo-striate and cortico-cortical projections

The superimposed maps from opposing hemifields reported for V1 in albinism are taken as macroscopic evidence for a cortical organization pattern termed “interleaved representation” which appears to be the only organization pattern available to primates with albinism (Guillery, 1986; Guillery et al., 1984). Here the former ocular dominance domains are reassigned to hemifield dominance domains to accommodate the abnormal input from the ipsilateral visual field in albinism. Importantly, this cortical representation can be explained by largely stable, geniculo-striate projections. In turn, independent visual functioning of the two hemifield representations is assumed to be due to adaptations of the intra-cortical micro-circuitry in V1 (Hoffmann and Dumoulin, 2015; Sinha and Meng, 2012). The propagation of this pattern beyond V1 indicates, as reported here and in previous studies (Hoffmann et al., 2003; Kaule et al., 2014), a largely stable cortico-cortical connectivity. This stability is further supported by our observation of a similar V1-sampling extent in V2 for controls and albinism. Remarkably, the V1-sampling extent appears altered beyond V2. In fact, while an unaltered gross-connectivity serves the propagation of the retinotopic maps through the hierarchy of the early visual cortex, the altered sampling of V3 from V1 might reflect a specific adaptation to the abnormal cortical input in albinism. This is suggested by the comparison of the mechanisms available to accommodate the ‘extra-map’, i.e. from the ipsilateral hemifield, in striate vs extrastriate cortex: At the level of extrastriate cortex, the accommodation of the representation of the ipsilateral visual field is much more demanding, since, at this stage, most neurons normally receive binocular input (Felleman and Van Essen, 1987; Kaule et al., 2014; Maunsell and van Essen, 1983; Tanabe et al., 2005). While in albinotic V1 the obsolete ocular dominance domains can be ‘simply’ reassigned to hemifield dominance domains, in extrastriate cortex no such spare resources appear to be available. Consequently, part of the neural resources normally allocated for processing the contralateral visual field must be made available for processing the additional input from the ipsilateral visual field. This is expected to result in an – at the mesoscopic scale – altered representation of the visual information from V2 onwards. As a result, the sampling by V3 is expected to be altered. Accordingly, our results for the V1-sampling extent in V3 might, therefore, reflect these extra-striate adaptations in albinism. Further studies are needed to elucidate this process and to identify the underlying adaptive mechanisms. Taken together, our findings highlight the dominance of conservative developmental mechanisms in human albinism, but at the same time indicate that plasticity shaping the input to V3 contributes to tuning the cortico-cortical connectivity to the altered visual input.

## 5. Conclusion

Albinism has a profound effect on the structure and function of the visual system, providing a compelling model to study the interplay of stability and plasticity in the human visual system. Our findings demonstrate the absence of extensive reorganization and gross stability of geniculo-striate and cortico-cortical projections. The adjustments of the cortico-cortical connections at the level of V3 might be critical to support independent processing of two opposing hemifields within the same hemisphere.

## Acknowledgement

We thank Prof. Serge O. Dumoulin, Dr. Koen Haak and Dr. Alessio Fracasso for the valuable discussions, and greatly appreciate the cooperation of the study participants. In addition, we thank the Center for Magnetic Resonance Research at the University of Minnesota, for providing the sequence files for multi-band EPI acquisition. This project was supported by European Union’s Horizon 2020 research and innovation programme under the Marie Sklodowska-Curie grant agreement (No. 641805) and the German research foundation (DFG, HO 2002/10-3) to M. B. H.

## Conflict of interests

None of the authors has potential conflicts of interest to be disclosed.

